# Castration-resistant prostate cancer: Androgen receptor inactivation induces telomere DNA damage, and damage response inhibition leads to cell death

**DOI:** 10.1101/516138

**Authors:** Vidyavathi Reddy, Asm Iskander, Clara Hwang, George Divine, Mani Menon, Evelyn R. Barrack, G. Prem-Veer Reddy, Sahn-Ho Kim

## Abstract

Telomere stability is important for cell viability, as cells with telomere DNA damage that is not repaired do not survive. We reported previously that androgen receptor (AR) antagonist induces telomere DNA damage in androgen-sensitive LNCaP prostate cancer cells; this triggers a DNA damage response (DDR) at telomeres that includes activation of ATM, and blocking ATM activation prevents telomere DNA repair and leads to cell death. Remarkably, AR antagonist induces telomere DNA damage and triggers ATM activation at telomeres also in 22Rv1 castration-resistant prostate cancer (CRPC) cells that are not growth inhibited by AR antagonist. Treatment with AR antagonist enzalutamide (ENZ) or ATM inhibitor (ATMi) by itself had no effect on growth in vitro or in vivo, but combined treatment with ENZ plus ATMi significantly inhibited cell survival in vitro and tumor growth in vivo. By inducing telomere DNA damage and activating a telomere DDR, an opportunity to inhibit DNA repair and promote cell death was created, even in CRPC cells. 22Rv1 cells express both full-length AR and AR splice variant AR-V7, but full-length AR was found to be the predominant form of AR associated with telomeres and required for telomere stability. Although 22Rv1 growth of untreated 22Rv1 cells appears to be driven by AR-V7, it is, ironically, expression of full-length AR that makes them sensitive to growth inhibition by combined treatment with ENZ plus ATMi. Notably, this combined treatment approach to induce telomere DNA damage and inhibit the DDR was effective in inducing cell death also in other CRPC cell lines (LNCaP/AR and C4-2B). Thus, the use of ENZ in combination with a DDR inhibitor, such as ATMi, may be effective in prolonging disease-free survival of patients with AR-positive metastatic CRPC, even those that co-express AR splice variant.

## Introduction

The critical role of the androgen receptor (AR) in prostate cancer cell proliferation and survival is the enduring basis for treating advanced prostate cancer with drugs that block AR function or androgen biosynthesis (1, 2). However, a relentless challenge is the development of resistance to these treatments, referred to as castration-resistant prostate cancer (CRPC) (3). Remarkably, CRPC still relies on AR (4, 5), indicating a need to more fully understand the role of AR in cell survival. In this regard, we have discovered a role of AR in prostate cancer cell telomere stability (6, 7). Notably, inactivation of this role of AR creates a DNA damage response (DDR) target, inactivation of which blocks repair and promotes cell death (8).

Telomeres are the DNA-protein structures that cap the ends of linear chromosomes, which are double-stranded DNA with a single-stranded overhang (9). Telomeres contain many different proteins that play a role in the maintenance of telomere stability; the best characterized are the six proteins (TRF1, TRF2, Rap1, TIN2, POT1 and TPP1) that comprise a complex known as shelterin (10). Shelterin and accessory proteins play a critical role in protecting chromosome ends from being recognized as lesions by the DNA damage machinery (11). Inhibition or down regulation of these proteins causes telomere dysfunction, a condition in which unprotected chromosome ends resemble damaged DNA and recruit DDR factors, such as 53BP1, Mre11, and phosphorylated (activated) forms of H2AX, ATM and Rad17 (12), which in turn trigger cell cycle checkpoint activation (11, 13). If damage can be repaired, the cell will remain viable; otherwise, cell death pathways will be activated (14). Thus, telomere stability is important for cell viability, and telomere DNA damage creates an opportunity to inhibit telomere DNA repair and activate cell death (8).

AR antagonists induce telomere DNA damage in AR-positive LNCaP prostate cancer cells, and a DDR that has the features of a bonafide telomere DDR, namely, activation of ATM, as indicated by an increase in phosphorylated ATM (pATM) at telomeres (6–8). Combined treatment with AR antagonist and ATM inhibitor (ATMi) increases the level of replication protein A (RPA, a marker of unrepaired single stranded DNA) at telomeres, indicating that repair of damaged telomere DNA has been blocked. This combined treatment increases the fraction of cells with sub-G_1_ DNA content (i.e., dead cells), presumably a result of cells entering mitosis with a level of telomere DNA damage that is incompatible with survival (8).

AR antagonist-induced telomere DNA damage in LNCaP prostate cancer cells appears to be mediated by telomere-associated AR, as AR-chromatin immunoprecipitate (AR-ChIP) contains telomeric DNA, isolated telomeric chromatin contains AR, and AR co-immunoprecipitates and colocalizes with shelterin proteins TIN2, TRF1 and TRF2 (6, 7). In addition, this telomere damage is independent of AR transcriptional activity, independent of p53 status, and not due to down-regulation of telomerase (6–8). Notably, AR antagonist does not cause genome-wide DNA damage, and agents such as etoposide that cause genome-wide DNA damage do not induce telomere DNA damage (7).

The AR antagonist bicalutamide induces telomere DNA damage in a variety of prostate cancer cells that express different forms of AR (7, 8): LNCaP cells that express mutant AR (8), LAPC4 cells that express wild-type AR (7), and 22Rv1 cells (15) that express both full-length AR (f-AR) and a constitutively active AR splice variant, AR-V7, that lacks the ligand-binding domain (8). The ability of AR antagonist to induce telomere DNA damage in CRPC 22Rv1 cells is intriguing because proliferation of these cells is ligand-independent and resistant to growth inhibition by AR antagonist.

Enzalutamide (ENZ) is a second-generation AR antagonist widely used to treat patients with CRPC (16), however, even tumors that initially respond eventually develop resistance (3). The 22Rv1 human CRPC cell line is resistant to growth inhibition by ENZ; thus it is a useful model to investigate therapeutic approaches to combat ENZ resistance. The AR splice variant AR-V7 accounts for androgen-independent growth and survival of 22Rv1 cells, as knockdown of AR-V7 with siRNA inhibits survival (15). The AR-V7 splice variant mediates ENZ resistance in 22Rv1 cells, is up-regulated during progression to CRPC in patients (15, 17), and is expressed in 19-59% of patients with AR-positive metastatic CRPC (18, 19).

Therefore, we investigated the role of full-length AR and splice variant AR-V7 in telomere stability and in the telomere DDR to AR antagonist ENZ in CRPC 22Rv1 cells. We also describe the growth inhibitory effect of combined inhibition of AR and DNA repair on CRPC 22Rv1 tumors in vivo.

## Materials and Methods

### Cell culture

LNCaP (ATCC), 22Rv1 (ATCC), C4-2B (MD Anderson Cancer Center) and LNCaP/AR (a gift from Drs. Robert Reiter and Charles Sawyers) cells were grown in RPMI (Gibco BRL) containing 10% fetal bovine serum (FBS), 2.5 mM glutamine, 100 μg/ml streptomycin and 100 U/ml penicillin (complete medium). Exponentially growing cells were treated as described in figure legends, in fetal calf serum (FCS)-containing medium. The concentration of AR antagonist that induces telomere DNA damage in prostate cancer cells is lower in charcoal-stripped fetal calf serum (CSS) than in untreated serum (FCS) (Fig. S1). However, to avoid confounding the effect of AR antagonist with the steroid hormone-depleting effect of CSS on AR activity, we use hormone-replete FCS in all experiments unless noted otherwise.

### Indirect immunofluorescence

The immunofluorescent staining of cells grown on glass slides was performed as described (20). Cells were fixed with 4% paraformaldehyde, permeabilized with 0.5 % Triton X-100 and incubated at 4°C overnight with rabbit polyclonal antibodies against TIN2 (20), γ-H2AX (i.e., phosphorylated-H2AX) (Upstate), or AR (AR-N20; Santa Cruz), or mouse monoclonal antibodies against AR (AR-414; Santa Cruz), AR-V7 (AG10008; Precision) or pATM (10H11-E12, which detects phosphorylation of ATM at serine 1981; Cell Signaling). Cells were then washed and stained with FITC-labeled goat-anti-rabbit-IgG and/or Texas Red-labeled goat-anti-mouse-IgG (Molecular probes) secondary antibodies. Images of cells were acquired on an LSM-410 confocal microscope (Zeiss). Labeled foci were counted in enlarged photographs.

### TIF response

Telomere DNA damage (telomere dysfunction)-mediated activation of DDR signaling leads to the phosphorylation of H2AX at telomeres. Therefore, cells containing immunofluorescent foci of phosphorylated H2AX (γH2AX) that colocalizes with telomeric protein TIN2, which are referred to as telomere-dysfunction induced *foci* (TIF), are scored as a measure of DDR, as described previously (6, 7, 12). Individual cells with >5 γ-H2AX foci were defined as having a TIF response, as few untreated cells have >5 TIF foci. Eighty cells/treatment were counted in each of three separate experiments. For example, 85% of untreated LNCaP cells have <5 TIF foci (ref. 6); by contrast, 85% of cells treated with AR antagonist have > 5 TIF foci/cell (28% have 6-10 foci/cell, 45% have 11-20 foci/cell, and 12% have >20 foci/cell (6).

### Cell extracts and Western blotting

Cells were harvested by trypsinization, washed with PBS and suspended in Buffer A (50 mM Tris-HCl, pH 7.4, 250 mM NaCl, 0.1% Triton X-100, 5 mM EDTA, 50 mM NaF, and 0.1 mM Na_3_VO_4_) supplemented with protease inhibitor mixture (P-8340, Sigma) at a density of 2×10^7^ cells/ml as described (21, 22). Cells were then subjected twice to 30 pulses of sonication with a Branson Sonifier 250 set at output control 2 and duty cycle 20, with intermittent cooling on ice. The sonicated cell extract was cleared by centrifugation in an Eppendorf centrifuge at 12,500 rpm for 10 min. For Western blotting, membranes were probed with antibodies against AR (AR-N20, Santa Cruz), AR-V7 (AG10008, Precision) or GAPDH (AB2302, Millipore). Immunoreactive bands were developed using horseradish peroxidase-conjugated secondary antibodies and SuperSignal WestPico chemiluminescent substrate (Pierce), and visualized using X-ray film.

### RT-PCR analysis

Total RNA was prepared as described (22). RNA was reverse transcribed using random hexamers and oligo (dT) primer and Transcriptor Reverse Transcriptase (Roche Applied Science) according to the manufacturer’s instructions. PCR of cDNA was carried out using Platinum PCR SuperMix (Invitrogen). PCR primers for AR were 5’-tcagttcacttttgacctgctaa (forward) and 5’-gtggaaatagatgggcttga (reverse), and for GAPDH were 5’-gagatccctccaaaatcaagtg (forward) and 5’ ccttccacgataccaaagttgt (reverse). Cycle parameters were 94°C for 2 min, 94°C for 30 sec, 55 °C for 30 sec and 68°C for 1 min. AR was amplified for 30 cycles and GAPDH for 25 cycles.

### Chromatin immunoprecipitation (ChIP) analysis

Cells in culture dishes were fixed with 1% formaldehyde in PBS at room temperature for 60 min, harvested by scraping, washed with PBS, and lysed in 1% SDS, 50 mM Tris-HCl, pH 8.0, 10 mM EDTA at a density of 10^7^ cells/ml. The lysate was then sonicated using a Branson Sonifier 250 under the following conditions: output control of 5.5 and duty cycle of 50% with six cycles of 20 sec with intermittent cooling on ice, and cleared by centrifugation in an Eppendorf centrifuge at 12,500 rpm for 10 min. The cleared lysate (0.2 ml) was diluted with 1.2 ml Buffer containing 0.01% SDS, 1.1% Triton X-100, 1.2 mM EDTA, 16.7 mM Tris-HCl, pH 8.0, and 150 mM NaCl, incubated at 4°C overnight with 5 μg antibody [IgG (Santa Cruz), AR N-20 (Santa Cruz), GR (Cell signaling), PR (Cell Signaling), RNA Pol II (Imgenex), or Rap1 (Bethyl Lab)], and the antibody-bound material was then precipitated with 30 μl protein-G Sepharose beads (Invitrogen) that had been pre-equilibrated with 30 μg bovine serum albumin (BSA) and 5 μg sheared *Escherichia coli* DNA for 30 min at 4°C. In order to isolate DNA from the ChIP pellet (ChIP DNA), cross-linking was reversed at 65°C for 4 h, treated with RNAase A and proteinase K at 37 °C, and extracted with 0.5 ml phenol/chloroform/isoamylalcohol. In order to probe ChIP DNA for AR binding sites in the PSA gene, NDR G1 gene and telomeres, ChIP DNA was subjected to PCR using primers for PSA ARE III: 5’- cttctagggtgaccagagcag (forward) and 5’-gcaggcatccttgcaagatg (reverse), for NDR-G1 ARE 5’- gccacctgggtagctttgta (forward) and 5’-agaggagccgccaaattaaa (reverse) and for chromosome 17p telomeres (Chr. 17p-Tel): 5’-gaatccacggattgctttgtgtactt (sub-telomeric forward) and 5’-tgctccgtgcatctggcatc(ccctaa)_5_ (telomeric reverse). Cycle parameters for ARE III were 94°C for 2 min, 94°C for 30 sec, 55°C for 30 sec and 68°C for 1 min for 30 cycles. Cycle parameters for telomeres were 95°C for 3 min 94°C for 30 sec, 60°C for 30 sec, 68°C for 1 min for 35 cycles.

### AR knockdown

Exponentially growing 22Rv1 cells (1.0 -2.0 x 10^5^ cells/well of a six-well plate) were transfected with 200 pmol siRNA targeting AR exon 1 (Ex1 siRNA, CAAGGAGGUUACACCAAA, to knock down both f-AR and AR-V7 (23)), AR exon 7 (Ex7 siRNA, UCAAGGAACUCGAUCGUAU, to knock down f-AR (23)), AR cryptic exon 3 (ExCE3 siRNA, GUAGUUGUGAGUAUCAUGA, to knock down AR-V7 (24)), or a control scrambled sequence (Santa Cruz), using Lipofectamine 2000 (Invitrogen) following the manufacturer’s instructions. Cryptic exon 3 (24) is also known as cryptic exon 3b (Welti et al., 2016 Eur Urol 70: 599-608). Cells were processed 36 hr later for immunofluorescence staining or Western blotting. In addition, transfected cells were treated with or without ATMi KU60019 (Selleck Chemicals, TX) for an additional 24 hr prior to colony formation assay.

### Colony formation assay

This procedure is essentially as described by Guzman et al (25). Cells (0.5-1.0 x 10^4^ cells/well of a six-well plate) were treated as described in figures for 24 hr, then washed to remove drugs and allowed to grow for 10-14 days, then fixed and stained with 0.01 % crystal violet (26). The areas of stained surviving cells in each plate were photographed and measured using the ImageJ program (25). The survival fraction was plotted relative to control (vehicle).

### 22Rv1 Xenografts

In order to test the effect of AR antagonist ENZ and ATM inhibitor KU59403 (Medkoo Bioscience, NC) on 22Rv1 tumors, athymic nude mice (Charles River) were inoculated subcutaneously with 4 × 10^6^ 22Rv1 cells as described by Wu et al.(27), and when tumor size reached about 200 mm^3^, tumor-bearing mice were randomly assigned to the following 4 treatment groups: vehicle (control, 6 mice), ENZ (enzalutamide, Selleckchem, TX) alone (7 mice); ATMi KU59403 alone (7 mice); ENZ + KU59403 (6 mice). All treatments were carried out 5 days/week for 4 weeks. KU59403 (25 mg/kg) was administered twice daily by i.p. injection (28). ENZ (50 mg/kg) was administered daily by oral gavage (29). Tumor growth was monitored by measuring tumor volume (30) twice a week. Treatments were carried out for 4 weeks, or until tumor volume reached 2,000 mm^3^ when mice were sacrificed and tumors were harvested.

### Immunohistochemistry

Harvested tumors were formalin-fixed, paraffin embedded, and five-micrometer sections cut to assess the activation of DDR signaling and induction of apoptosis in tumor cells. Activation of DDR signaling was assessed by immunohistochemical (IHC) staining of pATM using anti-pATM antibodies (Santa Cruz) and the IHC analysis Kit (Vector Laboratories), and apoptotic cell death was assessed by using a TUNEL assay Kit (InVitrogen), following the manufacturer’s suggested protocols.

### Statistics

Data are presented as mean +/- SD of three or more independent experiments. Statistical significance was calculated using the two-tailed Student t-test, using GraphPad Prism Software. A p-value <0.05 was considered significant.

## Results

### ENZ induces telomere damage in CRPC cells, and combining ENZ with an ATMi leads to cell death

Despite resistance to the growth inhibitory effect of ENZ, this AR antagonist nevertheless induces telomere DNA damage in 22Rv1 CRPC cells (Fig. 1A and Supplementary Fig. S2A), as well as in other human CRPC cell lines tested, namely, C4-2B (derived from an LNCaP xenograft propagated in castrated animals (31)) and LNCaP/AR (LNCaP cells engineered to overexpress AR (32)) (Fig. 1, B-C).

**Fig. 1:**
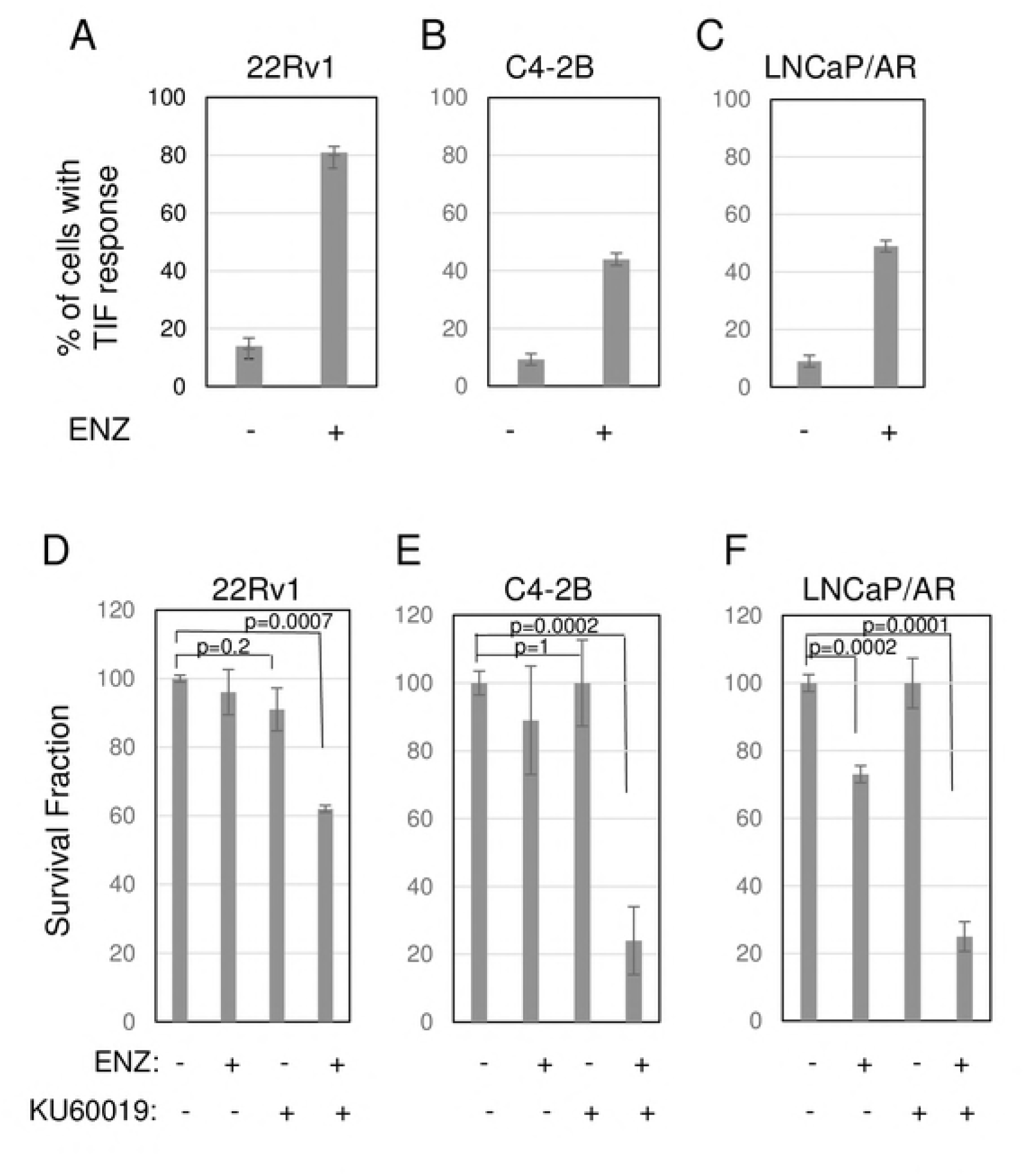
ENZ induces telomere damage, and inhibiting repair of damage with an ATMi leads to cell death, in multiple CRPC cell lines. A-C) ENZ induces telomere damage in CRPC cells. CRPC 22Rv1 (A), C4-2B (B) and LNCaP/AR (C) cells treated with ENZ were labeled with antibodies to DNA damage marker γ-H2AX and the telomere marker TIN2. Colocalization of γ-H2AX and TIN2 indicate DNA damage at telomeres. Cells with a TIF response to ENZ (>5 dual-labeled foci) were counted in enlarged (1000X) photomicrographs of representative fields. Data are expressed as mean ± SD of 3 independent experiments. **D-E) Combining ENZ with ATMi KU60019 leads to cell death**. 22Rv1 (D), C4-2B (E), LNCaP/AR (F) cells were treated with 5 μM ENZ in the presence or absence of 10 μM KU60019 for 24 hr, then washed to remove drugs and allowed to grow for 14 days (colony formation assay). The survival fraction is plotted relative to vehicle-treated controls; mean ± SD of 3 independent experiments.

ENZ also induces activation of ATM (Fig. S2B), a critical mediator of the telomere DDR (13). Notably, combined treatment of CRPC cells with ENZ (to induce telomere damage) plus the ATMi KU60019 (to inhibit the telomere DDR) leads to significant and substantial cell death in all three CRPC cell lines (Fig. 1, D-F). Thus, CRPC cells are vulnerable to treatments that target telomere stability and repair of telomere damage, though each alone has little or no effect on survival.

### Does AR splice variant AR-V7 play a role in telomere stability?

Our observation that AR antagonist induces telomere damage in AR-positive prostate cancer cells indicates a role of AR in telomere stability in these cells (6–8). Our studies using LNCaP cells indicate that this role is mediated by a subset of AR associated with telomeres (6). Although the AR splice variant AR-V7 cannot bind AR antagonist, AR-V7 might nonetheless play a role in telomere stability, for example, if it heterodimerized with full-length AR at telomeres. Therefore, we sought to determine whether AR-V7 is associated with telomeres in 22Rv1 cells.

One approach to identifying AR association with telomeres is dual-label immunofluorescence of AR that colocalizes with TIN2 in prostate cancer cells (6). The vast excess of AR relative to TIN2 presents a challenge. Therefore, prior to incubation with antibodies, we washed cells with cytoskeleton buffer (containing 0.1 M NaCl) plus 0.5% Triton X-100 to extract loosely bound cytoplasmic and nuclear proteins, as described by others (33, 34). This protocol extracted a lot of loosely bound nuclear AR, as indicated by a large decrease in nuclear AR staining, and made it much easier to identify colocalization of a subset of nuclear AR with TIN2 at telomeres (6). Nonetheless, even under these conditions, AR staining vastly exceeds TIN2 staining (Fig. 2A, vehicle panels).

**Fig. 2:**
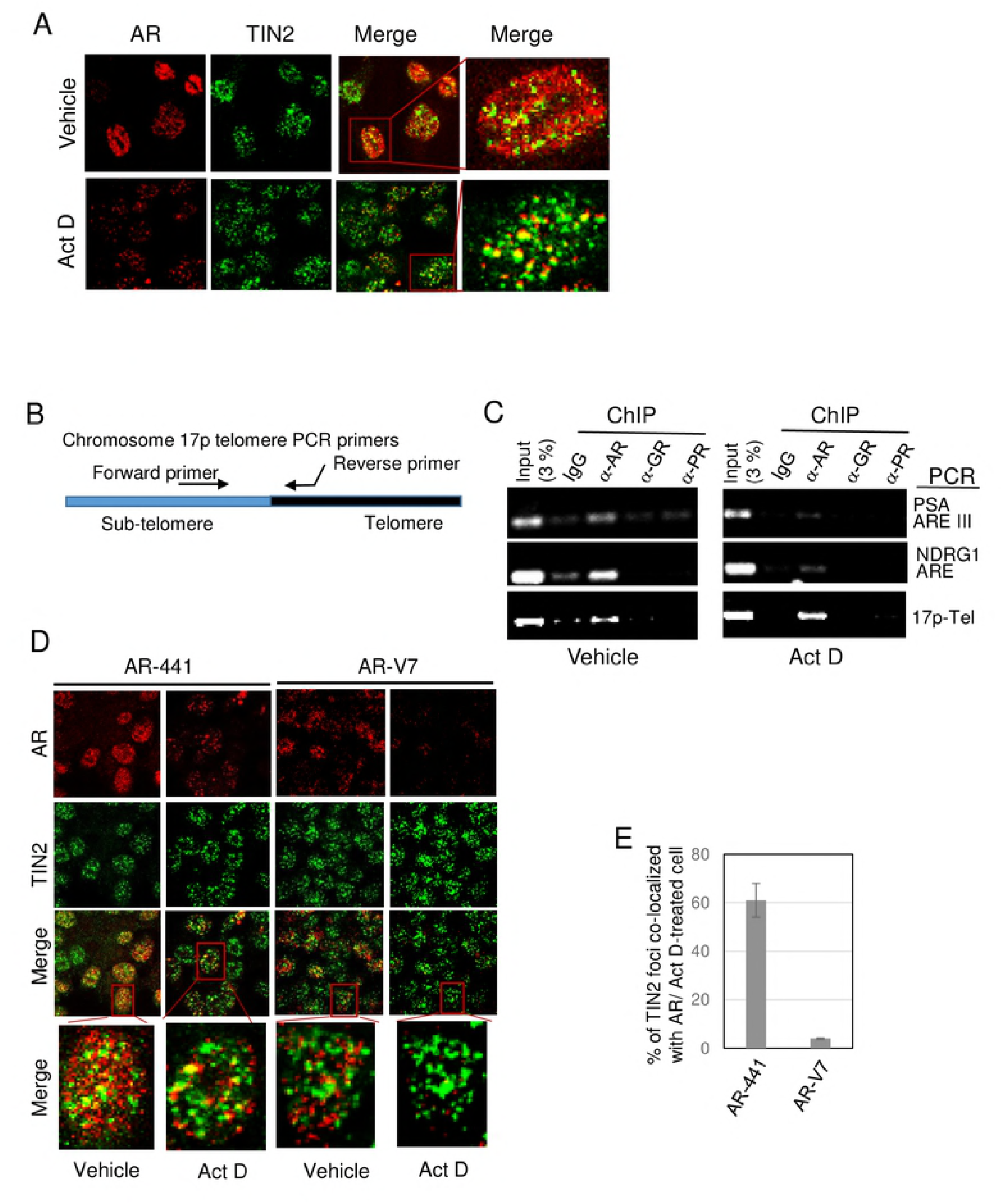
Telomere-associated AR in prostate cancer cells. A-D) Actinomycin D-resistant AR is preferentially associated with telomeres. **A) Actinomycin D treatment facilitates immunofluorescent identification of AR at telomeres.** LNCaP cells were treated with or without 0.5 μg/ml actinomycin D (Act D) for 4 hr, then fixed, permeabilized, and equilibrated in cytoskeleton buffer, and then subjected to dual labeling with antibodies against AR (AR-441, *red*) and TIN2 (*green*). Actinomycin D treatment greatly decreases the amount of nuclear AR without affecting TIN2. Dual-labeled foci (yellow in the *merge* panels) represent AR at telomeres, and are more evident in actinomycin D-treated cells. **B)** Most of the residual AR in Act D-treated cells in *A* was colocalized with TIN2 at telomeres. Foci in *A* were counted and data are expressed as the percentage of TIN2 foci colocalized with AR. Twenty cells were analyzed/treatment in each of 3 separate experiments. Error bars represent mean ± SD. **C)** Schematic of location of PCR primers used to amplify a region spanning the sub-telomere and telomere region of chromosome 17p. **D) Actinomycin D treatment decreases AR association with AREs but not with telomeres.** Chromatin was isolated from LNCaP cells that had been treated with or without actinomycin D and then fixed; chromatin immunoprecipitates (ChIPs) were prepared using antibodies against AR (N-20), GR, and PR, and analyzed by PCR for the presence of androgen response elements (AREs) of PSA and NDRG1 gene promoters, or chromosome 17p telomere DNA. **E-F) Full length AR, but not splice variant AR-V7, is associated with telomeres in 22Rv1 cells. E)** Immunofluorescent images of 22Rv1 cells that were treated with or without 0.5 μg/ml Act D for 4 hr, then labeled with TIN2 antibody and either antibody AR-441 (which recognizes both full-length AR and splice variant AR-V7) or an AR-V7-specific antibody are shown. The merge panels show colocalization of antibody AR441 with TIN2, but not of AR-V7 with TIN2 (both in vehicle-treated and actinomycin D-treated cells); this suggests that only full-length AR is associated with telomeres. **F)** Quantitation of labeled foci in actinomycin D-treated cells in **E.** Labeled foci were counted in twenty Act D-treated cells/group in each of three separate experiments, and expressed as the percentage of TIN2 foci colocalized with AR. Error bar represents mean ± SD, n=3.

We previously used actinomycin D to demonstrate that the role of AR in telomere stability is independent of AR transcriptional activity, as inhibiting the expression of AR-target genes in LNCaP cells with actinomycin D does not cause telomere DNA damage (7). This led us to hypothesize that there are two pools of nuclear AR protein in prostate cancer cells: one bound to chromatin where it functions as a transcription factor and its activity is sensitive to actinomycin D, and the other that is telomere-bound where it functions in maintaining telomere stability independent of AR transcriptional activity and is resistant to actinomycin D.

Notably, actinomycin D inhibits transcription as a result of its ability to intercalate into DNA; however, owing to differences in histone modifications and compactness of nucleosomes, actinomycin D intercalates and disrupts DNA-protein interactions at least three times more efficiently in euchromatin than in heterochromatin (35–37). Telomeric and subtelomeric chromatin is considered heterochromatin as it is enriched in epigenetic marks that are characteristic of heterochromatin, such as H3K9me3, H4K20me3, and hypoacetylated H3 and H4 (38). The heterochromatin state of telomeres is also evident from the presence of SIRT6, a histone deacetylase that promotes transcriptional silencing, and heterochromatin protein HP1-γ required for telomere cohesion (39, 40).

Thus, we hypothesized that actinomycin D might disrupt euchromatin-associated AR more efficiently than telomere-associated AR. We first tested this hypothesis using LNCaP cells that express only full-length AR. We treated cells with actinomycin D and then prepared them for immunolabeling of AR and TIN2. Actinomycin D treatment indeed decreased the amount of nuclear AR, but had no effect on TIN2 protein (Fig. 2A). Most notably, most of the cytoskeleton buffer-resistant AR in actinomycin D-treated cells was colocalized with TIN2 (Fig. 2A, Act D merge panels, and Fig. 2B), providing direct evidence for the presence of a subset of AR associated with telomeres. The presence of residual, actinomycin D-resistant AR at telomeres explains why actinomycin D treatment does not cause telomere DNA damage.

We further validated the association of actinomycin D-resistant AR with telomeres by analyzing AR-chromatin immunoprecipitate (AR-ChIP) for the presence of telomere DNA, using PCR primers that amplify sequence extending from the sub-telomere region of chromosome 17p into its telomere (17p-Tel) (Fig. 2C). As expected, AR-ChIP prepared from untreated LNCaP cells (Fig. 2D, vehicle) contains androgen response elements (AREs) of AR-target genes PSA and NDRG1. Notably, AR-ChIP also contains telomere DNA of chromosome 17p (Fig. 2D, vehicle), consistent with our previous finding of telomere repeat DNA in AR-ChIP of LNCaP cells (7).

By contrast, AR-ChIP prepared from actinomycin D-treated cells (Fig. 2D, actinomycin D) had a decreased presence of AREs; this is presumably due to the ability of actinomycin D to disrupt AR binding to euchromatin and inhibit transcription. Notably, however, actinomycin D had no effect on the presence of 17p-telomere DNA in AR-ChIP (Fig. 2D), affirming the association of actinomycin D-resistant AR with telomeres.

We next used actinomycin D to address the question whether the AR splice variant AR-V7 in 22Rv1 cells is associated with telomeres and whether AR-V7 plays a role in telomere stability. Dual immunofluorescence labeling of 22Rv1 cells was carried out using TIN2 antibody and either antibody AR-441 that recognizes both full-length AR and variant AR-V7, or antibody specific to AR-V7 (Fig. 2E). Colocalization of the N-terminal domain of AR (using antibody AR-441) with TIN2 at telomeres was evident in vehicle-treated cells, but AR-V7 was rarely seen at telomeres (vehicle panels in Figs. 2E, 2F); this suggests that it is predominantly full-length AR associated with telomeres. Treatment of 22Rv1 cells with actinomycin D and subsequent washing with cytoskeleton buffer plus 0.5% Triton X-100 allowed for a large decrease in nuclear AR staining with no change in telomere TIN2 (Fig. 2E, compare vehicle vs. Act D). The predominant colocalization of AR-441 with TIN2, and relative lack of colocalization of AR-V7 with TIN2, were evident whether or not cells were pre-treated with actinomycin D (Fig. 2E), but identification and quantitation of colocalized foci was facilitated by the removal of soluble and loosely bound AR (Fig. 2F).

The apparent presence of a small subset of AR-V7 at telomeres in 22Rv1 cells (Fig. 2F) suggests a possible role in telomere stability, perhaps via heterodimerization with full-length AR (41). To address this question, we tested the effect of knockdown of f-AR and/or AR-V7 on telomere stability in 22Rv1 cells. We transfected cells with siRNA targeting (a) exon 1 to knockdown both f-AR and AR-V7, (b) exon 7 to knockdown only f-AR or (c) cryptic exon 3 (CE3) to knockdown only AR-V7 (Fig. 3A) (15). Knockdown of both f-AR and AR-V7 was slightly more effective than knockdown of only f-AR (p= 0.032) (Fig. 3C); since cells with only f-AR knockdown still express AR-V7 (Fig. 3A), a small contribution of AR-V7 to telomere stability cannot be ruled out. Knockdown of only AR-V7 caused only a low level of telomere DNA damage (Fig. 3C), suggesting a small contribution of AR-V7 to telomere stability; however, we cannot rule out the possibility that this low level of telomere dysfunction was caused by the partial decrease in f-AR in these cells (Fig. 3A). Thus, it appears that full-length AR is the predominant form of AR at telomeres (Fig. 2, E-F), and the predominant form of AR required for telomere stability (Fig. 3) in 22Rv1 cells.

**Fig. 3:**
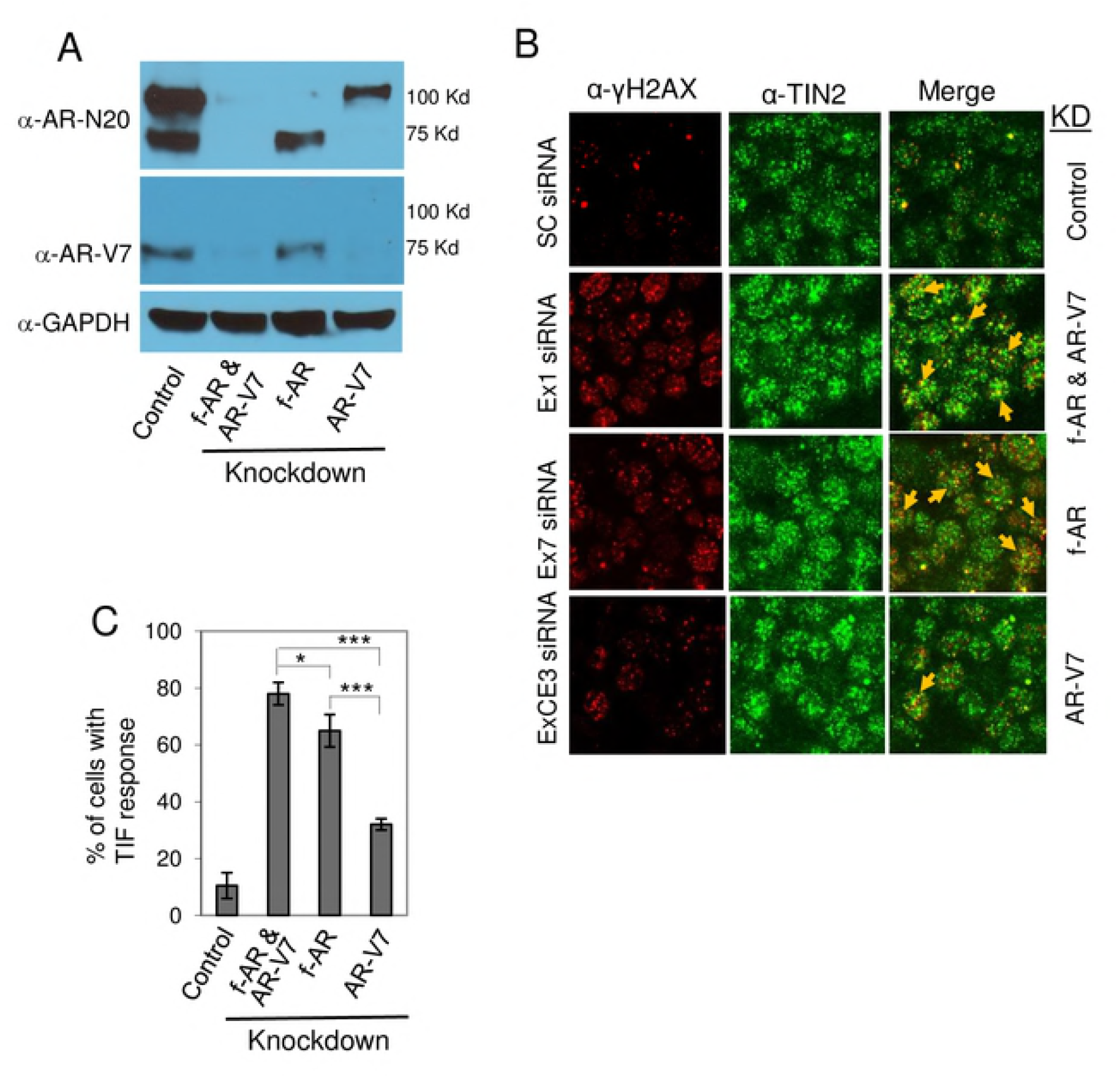
Effect of knockdown of full-lengh AR or AR-V7 on telomere DNA damage in 22Rv1 cells. 22Rv1 cells were transfected with siRNA targeting AR exon 1 to knockdown both full-lengh AR (f-AR) and AR-V7, AR exon 7 to knockdown only f-AR, or AR exon CE3 to knockdown only AR-V7. Scrambled siRNA was used as control. **A)** Cell extracts were prepared, and western blot analysis was performed using an AR antibody (N-20) that recognizes both f-AR and AR-V7, an AR-V7-specific antibody, or a GAPDH antibody. **B)** The effect of knockdown of f-AR and AR-V7, or of only AR-V7, on telomere DNA damage was assessed by dual immunofluorescent staining of the DNA damage marker γ-H2AX (red) and the telomere marker TIN2 (green). Colocalization of γ-H2AX and TIN2 (indicated by yellow arrows) is shown in the ‘merge’ panels. **C)** Bar chart of the percentage of cells with a TIF response, a measure of DNA damage at telomeres. Eighty cells/treatment were counted in each of three separate experiments; mean ± SD, n=3. *, p=0.04; ***, p=0.0001.

We next investigated the role of full-length AR and AR splice variant AR-V7 in the cell death response to ATMi. Knockdown of full-length AR alone did not affect 22Rv1 cell survival, presumably because AR-V7 was still expressed; however, knockdown of full-length AR plus treatment with ATMi KU60019 significantly decreased cell survival (p = 0.001) (Fig. 4), similar to the effect of AR antagonist ENZ plus ATMi (Fig.1D). Knockdown of AR-V7 alone did not decrease survival in hormone-replete FCS-containing medium (Fig. 4) (see Materials and Methods, and Fig. S1); this is consistent with the observation that knockdown of AR-V7 decreases survival only in hormone-depleted charcoal-stripped serum (CSS)-containing medium (15). According to Dehm and colleagues (15), knockdown of AR-V7 in the presence of androgen restores androgen responsiveness; with AR-V7 knocked down, 22Rv1 cells grow in response to androgen and this effect is blocked by AR antagonist, presumably via full-length AR. Thus, inactivation of full-length AR, when AR-V7 is knocked down, decreases survival of 22Rv1 cells (15); this is similar to the effect of knockdown of full-length AR other CRPC cells that express only full-length AR and rely on AR for survival (4, 5). This likely explains why knockdown of both full-length AR and variant AR-V7 decreased survival and why survival was not further reduced by additional treatment with ATMi (Fig. 4). As siRNA is not yet a reliable method for treatment in vivo, therefore we next tested the effect of combined antagonism of full-length AR plus inhibition of ATM on CRPC tumor growth in vivo.

**Fig. 4:**
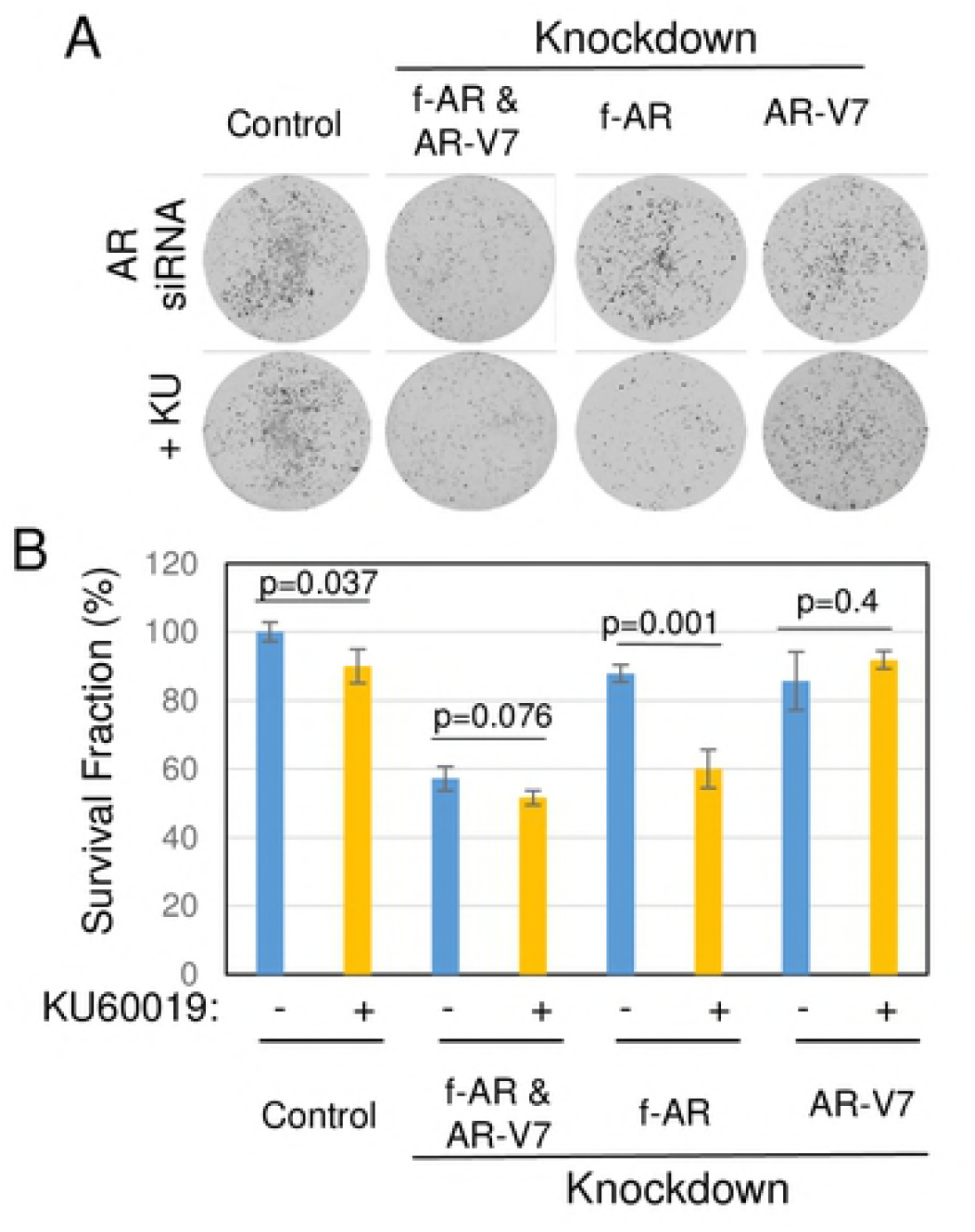
Effect of f-AR or AR-V7 knockdown in the presence of ATMi in 22Rv1 cells. **A)** 22Rv1 cells were transfected with siRNAs targeting AR exon 1 to knockdown both f-AR and AR-V7, AR exon 7 to knockdown only f-AR, or AR exon CE3 to knockdown only AR-V7. Scrambled siRNA was used as control. Transfected cells were then treated for 24 hr with or without 10 μM ATMi KU60019, then washed to remove drugs and subjected to a colony formation assay (14 day growth). B) Bar chart shows the survival fraction of cells from the colony formation assay, expressed relative to the survival of control cells transfected with scrambled siRNA and treated with vehicle; mean ± SD of 3 independent experiments.

### Treatment with ENZ plus ATMi suppresses CRPC 22Rv1 tumor growth in vivo

We treated established CRPC 22Rv1 tumors with vehicle, ENZ alone, ATMi alone, or ENZ in combination with ATMi (Fig. 4). We used the ATMi KU59403, which has been shown previously to have favorable pharmacokinetic properties and bioavailability in nude mice (28). Importantly, when administered by itself, KU59403 has no cytotoxic effects on vital organs or on body weight of mice (28).

Combined treatment of CRPC 22Rv1 tumor-bearing mice with ENZ plus ATMi dramatically inhibited tumor growth compared to the other treatments (Fig. 5A; individual tumor growth curves are shown in Fig. S3B. Not surprisingly, ENZ alone or ATMi alone had no significant effect on tumor growth (Fig. 5A and Fig. S3). Plotting the log of tumor volume vs. time allowed us to calculate tumor doubling times, which showed that only combined treatment with ENZ plus ATMi KU59403 slowed tumor growth (Fig. 5B). Notably, of 6 mice in the combined treatment group, tumor in 1 mouse became undetectable by day 9, and tumors in 2 other mice started to decrease in size beginning at day 20 (Fig. S3B). Kaplan-Meier analysis revealed a significant survival benefit in the combined treatment group (Fig. S4 and Table S1). Also, consistent with earlier reports (28), ATMi KU59403 was safe *in vivo* as it had no detrimental effect on body weight when administered alone or in combination with ENZ, compared to controls (Fig. 5C).

**Fig. 5:**
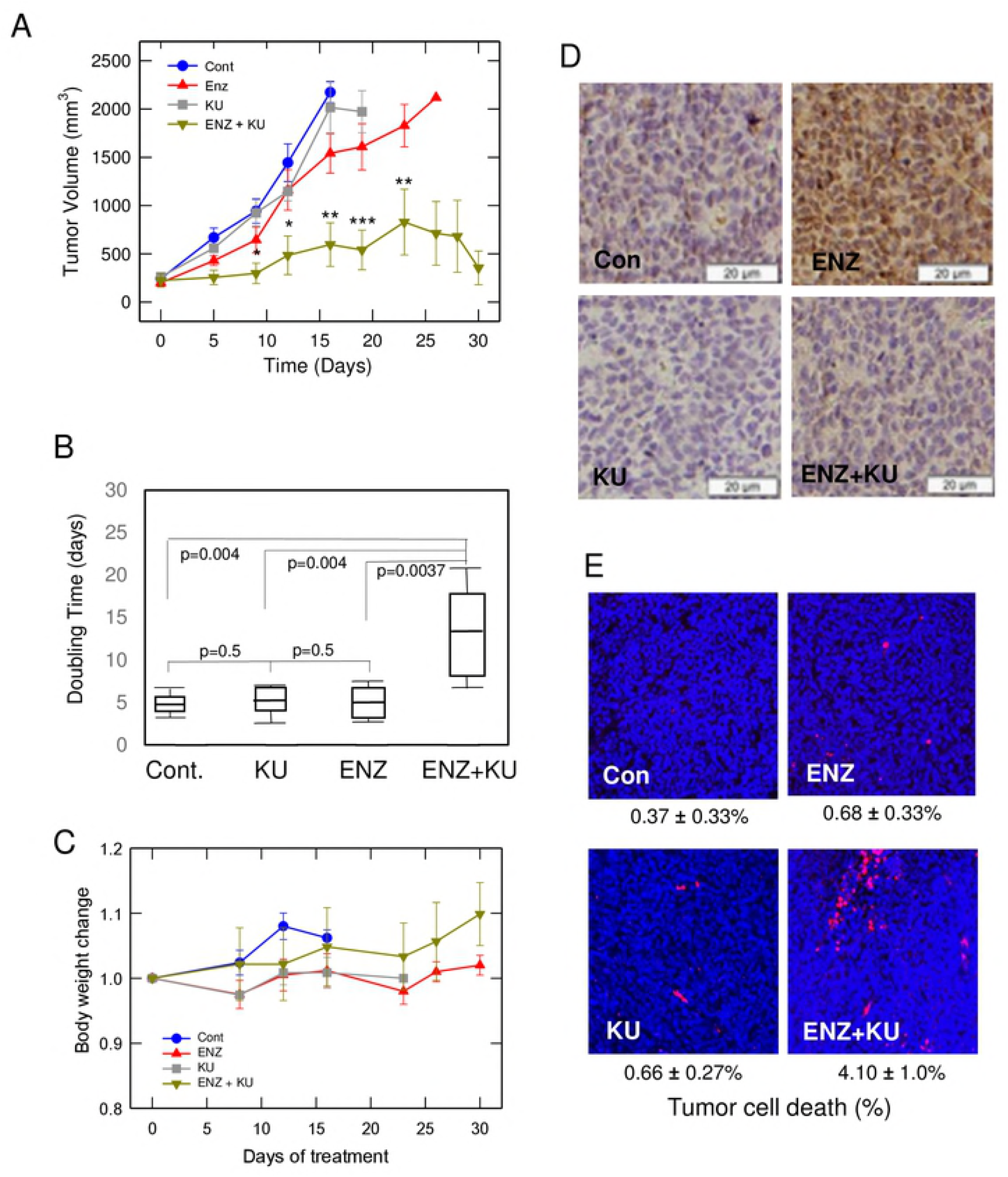
Combined treatment with AR antagonist ENZ plus ATMi inhibits growth of CRPC 22Rv1 xenograft tumors in mice that are resistant to each drug alone. 22Rv1 tumor-bearing athymic nude mice were randomly assigned to vehicle control (Cont), ENZ, ATMi KU59403 (KU), or combined ENZ+KU treatment for 4 weeks. **A)** Tumor size over time is presented as mean tumor volume (mm^3^) of each treatment group (n= 6 or 7 mice/group). Error bars represent standard deviation. Statistical analysis was performed for comparison between ENZ alone or KU alone and KU+ENZ treatments: *, p<0.05; **, p<0.001; ***, p<0.0001. **B)** The doubling time of each tumor was calculated from a plot of log tumor volume vs. time. Doubling times of each treatment group are presented as Box-Whisker plots; horizontal lines represent mean, first and third quartiles, and whiskers represent the minimum and maximum doubling time of each group. The doubling time (days, mean + SEM) of each group was: Control, 4.94 + 1.0; KU, 5.50 + 1.36; ENZ, 5.46 + 1.36; ENZ+KU, 12.71 + 4.38. P values are shown in the chart. **C)** Body weight of mice during the treatment period, relative to day 0 of treatment of each group. **D)** ENZ induces ATM activation in 22Rv1 xenograft tumors. Immunostaining of pATM is shown in a representative 22Rv1 xenograft tumor section from each treatment group. **E)** Evaluation of cell death in serial sections of 22Rv1 tumors. TUNEL assay to detect cell death was performed as described in the manufacturer’s protocol (InVitrogen). Images show ~1, 000 cells (blue) in a representative tumor tissue section from each treatment group. Cell death was analyzed by counting the percentage of cells that were dead (red) in each image.

Notably, treatment of 22Rv1 tumor-bearing mice with ENZ alone activated ATM (pATM) in vivo (Fig. 5D), as it did in vitro (Fig. S2B), but as expected did not cause cell death (Fig. 5E, TUNEL assay). Combined treatment with ENZ plus ATMi suppressed tumor growth (Figs. 5A and S3) and increased cell death (Fig. 5E), presumably a result of ATMi blocking DNA repair in cells with a DDR and activated pATM, so that cells with too much damage undergo cell death (8, 14, 42).

## Discussion

The expression of constitutively active AR splice variants such as AR-V7, which lacks the ligand binding domain, is considered an important mechanism of AR antagonist-resistant growth of metastatic CRPC (15, 17). However, because full-length AR is expressed in 80-100% of AR-positive CRPC metastases (18, 43), and because full-length AR, not AR-V7, is associated with telomeres and is critical for telomere stability, CRPC metastases are expected to be vulnerable to telomere DNA damage by AR antagonist, and activation of a telomere DDR that can be inhibited by an ATMi to promote cell death, as demonstrated in CRPC 22Rv1 cells and tumors, and in CRPC cell lines C4-2B and LNCaP/AR (this study). Thus, our data suggest that the use of ENZ in combination with a DDR inhibitor, such as an ATMi, may be effective in prolonging disease-free survival of patients with AR-positive mCRPC, even the 19-59% that co-express an AR splice variant (18, 43).

The AR is a well-characterized transcription factor that regulates the expression of many genes (19), including many that play a role in DNA repair (44, 45). In those studies, the growth inhibitory effect of ionizing radiation-induced genome-wide DNA damage was enhanced by AR inactivation, and this enhancement was inferred to be the result of decreased expression of DNA repair genes (44, 45). By contrast, our data indicate that AR antagonist induces telomere DNA damage independent of an effect on AR transcriptional activity (7). In addition, AR inactivation does not cause genome-wide DNA damage (6, 44), but by inducing telomere DNA damage and activating a telomere DDR, an opportunity to inhibit DNA repair is created, the consequence of which is to promote cell death, as genome integrity is required for survival. When applied to the treatment of CRPC 22Rv1 cells and tumors, combined treatment with AR antagonist plus ATMi significantly inhibited growth, whereas each drug alone was ineffective. This feature fits the criterion for synthetic lethality (46), although incomplete tumor elimination in our study suggests the need to target additional pathways or other components of the telomere DDR and repair pathway.

There is growing interest in combined targeting of AR and DNA repair in CRPC, using ENZ or androgen ablation plus an inhibitor of poly (ADP-ribose) polymerase (PARP) (47, 48) or an inhibitor of both Chk1 and Chk2 (49). Notably, these other studies focus on inhibiting genome-wide DNA repair, whereas our own studies focus on inhibiting the repair of telomere DNA damage. Cesare et al (50) have reported that the telomere DDR is functionally distinct from the genomic DDR.

Many cell types, including prostate cancer, use AR to modulate specific gene expression, but prostate cancer cells uniquely require AR to regulate cell proliferation and cell survival. Similarly, most cells do not need AR for telomere stability, but our studies have shown a critical role of AR in telomere stability in prostate cancer cells (6). Our data indicate that this role is mediated by a subset of AR associated with telomeres ((6) and this study), although it is not yet clear how AR is tethered to telomeres. Immunofluorescent colocalization of AR with telomere protein TIN2 was seen clearly after pretreating cells with actinomycin D and washing with cytoskeleton buffer to extract soluble nuclear AR (Fig. 2). Thus, actinomycin D treatment may be a useful tool for studying telomere-associated AR.

In summary, we have demonstrated that CRPC 22Rv1 tumor growth is resistant to AR antagonist or ATMi as monotherapies, but is significantly inhibited by combined treatment. CRPC cells that express both full-length AR and splice variant AR-V7 and are not growth inhibited by AR antagonist are inferred to depend on, and are said to be driven by AR-V7; but, ironically, it is their expression of full-length AR that makes them sensitive to growth inhibition by combined treatment with ENZ plus ATMi.

## Supporting information

**Fig. S1. The concentration of AR antagonist that induces telomere DNA damage in prostate cancer cells is lower in charcoal-stripped serum (CSS) than in untreated serum (FCS)**. Exponentially growing LNCaP cells were treated as indicated for 24 hr, in either FCS-containing medium (hormone-replete) or CSS-containing medium (hormone-depleted). To prepare cells for treatment with AR antagonist in CSS medium, exponentially growing LNCaP cells in FCS medium were washed twice with phenol red-free RPMI medium (Thermo Fisher Scientific) for 1 hr, and incubated in phenol red-free RPMI medium supplemented with 10% CSS (InVitrogen) for 26 hr prior to treatment with AR antagonists. After 24-hr treatment with AR antagonist, cells were labeled with antibodies to γ-H2AX (marker of DNA damage) and TIN2 (telomere specific protein), and cells with a TIF response (>5 dual-labeled foci/cell) were counted. Data are expressed as mean ± SD of 3 independent experiments. The concentration of bicalutamide (Casodex; from LKT Laboratories, MN) or the more potent ENZ that induces telomere DNA damage in LNCaP cells was lower in CSS (5 μM or 1 μM, respectively) than in untreated serum (FCS) (50 μM or 5 μM, respectively).

**Fig. S2. ENZ induces telomere damage (A) and activates ATM at telomeres (B) in CRPC cells. A)** 22Rv1 cells were treated without (control, Con) or with 5 μM ENZ in FCS-containing medium for 6 hr, then labeled with antibodies to DNA damage marker γ-H2AX (red) and the telomere marker TIN2 (green). Dual-labeled foci (indicated by yellow arrows) are shown in the ‘merge’ panel, indicating DNA damage at telomeres of ENZ-treated 22Rv1 cells. **B)** 22Rv1 cells were treated with or without 5 μM ENZ for 6 hr, then labeled with antibodies to phosphorylated ATM (pATM, red) and TIN2 (green). Colocalization of pATM (activated ATM) and TIN2 is shown in the ‘merge’ panels, indicating the presence of activated ATM at telomeres of ENZ-treated 22Rv1 cells.

**Fig. S3. Combined treatment with AR antagonist plus ATMi inhibits growth of CRPC 22Rv1 xenograft tumors in mice that are resistant to each drug alone.** These data supplement the data shown in Fig. 5. In this Figure, tumor volumes were normalized to the start of treatment on day 0, and are shown as fold change. **A)** Data for each group are shown as mean + SEM. *, p<0.05; **, p<0.001; ***, p<0.0001. **B)** Growth curves are shown for each tumor.

**Fig. S4. Kaplan-Meier survival analysis of 22Rv1 xenograft mice treated with AR antagonist plus ATMi**. Survival was defined as the number of days until sacrifice, when tumor size was ~2,000 mm^3^. Time to sacrifice was not adjusted for differences in tumor size at the start of treatment.

**Table S1:** Median Days to Sacrifice (tumor volume ~2000 mm^3^) For each treatment group, the median and 95% confidence interval for time to tumor volume ~2000 mm^3^ and log rank p-values were computed using the SAS procedure PROC LOGISTIC. Time to sacrifice was not adjusted for differences in tumor size at the start of treatment.

| Treatment | Median (95% CI) | Log rank p-value |  |  |
| --- | --- | --- | --- | --- |
|  |  | vs Control | vs ENZ | vs KU |
| Control | 16 (12, 16) |  |  |  |
| ENZ | 19 (12, 23) | 0.882 |  |  |
| KU59403 (KU) | 16 ( 9, 19) | 0.043 | 0.496 |  |
| ENZ+KU | >30 (23, >30) | <0.001 | 0.001 | <0.001 |

## References

1. de Bono JS, Logothetis CJ, Molina A, Fizazi K, North S, Chu L, et al. Abiraterone and increased survival in metastatic prostate cancer. The New England journal of medicine. 2011;364(21):1995–2005.

2. Scher HI, Fizazi K, Saad F, Taplin ME, Sternberg CN, Miller K, et al. Increased survival with enzalutamide in prostate cancer after chemotherapy. The New England journal of medicine. 2012;367(13):1187–97.

3. Feldman BJ, Feldman D. The development of androgen-independent prostate cancer. Nature reviews Cancer. 2001;1(1):34–45.

4. Haag P, Bektic J, Bartsch G, Klocker H, Eder IE. Androgen receptor down regulation by small interference RNA induces cell growth inhibition in androgen sensitive as well as in androgen independent prostate cancer cells. J Steroid Biochem Mol Biol. 2005;96(3-4):251–8.

5. Snoek R, Cheng H, Margiotti K, Wafa LA, Wong CA, Wong EC, et al. In vivo knockdown of the androgen receptor results in growth inhibition and regression of well-established, castration-resistant prostate tumors. Clinical cancer research: an official journal of the American Association for Cancer Research. 2009;15(1):39–47.

6. Kim SH, Richardson M, Chinnakannu K, Bai VU, Menon M, Barrack ER, et al. Androgen receptor interacts with telomeric proteins in prostate cancer cells. The Journal of biological chemistry. 2010;285(14):10472–6.

7. Zhou J, Richardson M, Reddy V, Menon M, Barrack ER, Reddy GP, et al. Structural and functional association of androgen receptor with telomeres in prostate cancer cells. Aging. 2013;5(1):3–17.

8. Reddy V, Wu M, Ciavattone N, McKenty N, Menon M, Barrack ER, et al. ATM Inhibition Potentiates Death of Androgen Receptor-Inactivated Prostate Cancer Cells with Telomere Dysfunction. The Journal of biological chemistry. 2015;290:25522–33.

9. Makarov VL, Hirose Y, Langmore JP. Long G tails at both ends of human chromosomes suggest a C strand degradation mechanism for telomere shortening. Cell. 1997;88(5):657–66.

10. de Lange T. Shelterin: the protein complex that shapes and safeguards human telomeres. Genes & development. 2005;19(18):2100–10.

11. Palm W, de Lange T. How shelterin protects mammalian telomeres. Annu Rev Genet. 2008;42:301–34.

12. Takai H, Smogorzewska A, de Lange T. DNA damage foci at dysfunctional telomeres. Curr Biol. 2003;13(17):1549–56.

13. Denchi EL, de Lange T. Protection of telomeres through independent control of ATM and ATR by TRF2 and POT1. Nature. 2007;448(7157):1068–71.

14. Smith J, Tho LM, Xu N, Gillespie DA. The ATM-Chk2 and ATR-Chk1 pathways in DNA damage signaling and cancer. Advances in cancer research. 2010;108:73–112.

15. Li Y, Chan SC, Brand LJ, Hwang TH, Silverstein KA, Dehm SM. Androgen receptor splice variants mediate enzalutamide resistance in castration-resistant prostate cancer cell lines. Cancer research. 2013;73(2):483–9.

16. Tran C, Ouk S, Clegg NJ, Chen Y, Watson PA, Arora V, et al. Development of a second-generation antiandrogen for treatment of advanced prostate cancer. Science. 2009;324(5928):787–90.

17. Guo Z, Yang X, Sun F, Jiang R, Linn DE, Chen H, et al. A novel androgen receptor splice variant is up-regulated during prostate cancer progression and promotes androgen depletion-resistant growth. Cancer research. 2009;69(6):2305–13.

18. Antonarakis ES, Lu C, Wang H, Luber B, Nakazawa M, Roeser JC, et al. AR-V7 and resistance to enzalutamide and abiraterone in prostate cancer. The New England journal of medicine. 2014;371(11):1028–38.

19. Pomerantz MM, Li F, Takeda DY, Lenci R, Chonkar A, Chabot M, et al. The androgen receptor cistrome is extensively reprogrammed in human prostate tumorigenesis. Nature genetics. 2015;47(11):1346–51.

20. Kim SH, Kaminker P, Campisi J. TIN2, a new regulator of telomere length in human cells [see comments]. Nature genetics. 1999;23(4):405–12.

21. Pelley RP, Chinnakannu K, Murthy S, Strickland FM, Menon M, Dou QP, et al. Calmodulin-androgen receptor (AR) interaction: calcium-dependent, calpain-mediated breakdown of AR in LNCaP prostate cancer cells. Cancer research. 2006;66(24):11754–62.

22. Bai VU, Cifuentes E, Menon M, Barrack ER, Reddy GP. Androgen receptor regulates Cdc6 in synchronized LNCaP cells progressing from G1 to S phase. Journal of cellular physiology. 2005;204(2):381–7.

23. Dehm SM, Schmidt LJ, Heemers HV, Vessella RL, Tindall DJ. Splicing of a novel androgen receptor exon generates a constitutively active androgen receptor that mediates prostate cancer therapy resistance. Cancer research. 2008;68(13):5469–77.

24. Hu R, Dunn TA, Wei S, Isharwal S, Veltri RW, Humphreys E, et al. Ligand-independent androgen receptor variants derived from splicing of cryptic exons signify hormone-refractory prostate cancer. Cancer research. 2009;69(1):16–22.

25. Guzman C, Bagga M, Kaur A, Westermarck J, Abankwa D. ColonyArea: an ImageJ plugin to automatically quantify colony formation in clonogenic assays. PloS one. 2014;9(3):e92444.

26. Golding SE, Rosenberg E, Valerie N, Hussaini I, Frigerio M, Cockcroft XF, et al. Improved ATM kinase inhibitor KU-60019 radiosensitizes glioma cells, compromises insulin, AKT and ERK prosurvival signaling, and inhibits migration and invasion. Molecular cancer therapeutics. 2009;8(10):2894–902.

27. Wu M, Kim SH, Datta I, Levin A, Dyson G, Li J, et al. Hydrazinobenzoylcurcumin inhibits androgen receptor activity and growth of castration-resistant prostate cancer in mice. Oncotarget. 2015;6(8):6136–50.

28. Batey MA, Zhao Y, Kyle S, Richardson C, Slade A, Martin NM, et al. Preclinical evaluation of a novel ATM inhibitor, KU59403, in vitro and in vivo in p53 functional and dysfunctional models of human cancer. Molecular cancer therapeutics. 2013;12(6):959–67.

29. Fokas E, Prevo R, Pollard JR, Reaper PM, Charlton PA, Cornelissen B, et al. Targeting ATR in vivo using the novel inhibitor VE-822 results in selective sensitization of pancreatic tumors to radiation. Cell death & disease. 2012;3:e441.

30. Jin RJ, Wang Y, Masumori N, Ishii K, Tsukamoto T, Shappell SB, et al. NE-10 neuroendocrine cancer promotes the LNCaP xenograft growth in castrated mice. Cancer research. 2004;64(15):5489–95.

31. Thalmann GN, Anezinis PE, Chang SM, Zhau HE, Kim EE, Hopwood VL, et al. Androgen-independent cancer progression and bone metastasis in the LNCaP model of human prostate cancer. Cancer research. 1994;54(10):2577–81.

32. Chen CD, Welsbie DS, Tran C, Baek SH, Chen R, Vessella R, et al. Molecular determinants of resistance to antiandrogen therapy. Nature medicine. 2004;10(1):33–9.

33. Nickerson JA, Krockmalnic G, Wan KM, Penman S. The nuclear matrix revealed by eluting chromatin from a cross-linked nucleus. Proceedings of the National Academy of Sciences of the United States of America. 1997;94(9):4446–50.

34. Compton DA, Yen TJ, Cleveland DW. Identification of novel centromere/kinetochore-associated proteins using monoclonal antibodies generated against human mitotic chromosome scaffolds. The Journal of cell biology. 1991;112(6):1083–97.

35. Berlowitz L. Chromosomal inactivation and reactivation in mealy bugs. Genetics. 1974;78(1):311–22.

36. Xu F, Zhang Q, Zhang K, Xie W, Grunstein M. Sir2 deacetylates histone H3 lysine 56 to regulate telomeric heterochromatin structure in yeast. Molecular cell. 2007;27(6):890–900.

37. Galati A, Micheli E, Cacchione S. Chromatin structure in telomere dynamics. Front Oncol. 2013;3:46.

38. Blasco MA. The epigenetic regulation of mammalian telomeres. Nat Rev Genet. 2007;8(4):299–309.

39. Canudas S, Houghtaling BR, Bhanot M, Sasa G, Savage SA, Bertuch AA, et al. A role for heterochromatin protein 1gamma at human telomeres. Genes & development. 2011;25(17):1807–19.

40. Imai S, Armstrong CM, Kaeberlein M, Guarente L. Transcriptional silencing and longevity protein Sir2 is an NAD-dependent histone deacetylase. Nature. 2000;403(6771):795–800.

41. Watson PA, Chen YF, Balbas MD, Wongvipat J, Socci ND, Viale A, et al. Constitutively active androgen receptor splice variants expressed in castration-resistant prostate cancer require full-length androgen receptor. Proceedings of the National Academy of Sciences of the United States of America. 2010;107(39):16759–65.

42. Helleday T, Petermann E, Lundin C, Hodgson B, Sharma RA. DNA repair pathways as targets for cancer therapy. Nature reviews Cancer. 2008;8(3):193–204.

43. Sun S, Sprenger CC, Vessella RL, Haugk K, Soriano K, Mostaghel EA, et al. Castration resistance in human prostate cancer is conferred by a frequently occurring androgen receptor splice variant. The Journal of clinical investigation. 2010;120(8):2715–30.

44. Goodwin JF, Schiewer MJ, Dean JL, Schrecengost RS, de Leeuw R, Han S, et al. A hormone-DNA repair circuit governs the response to genotoxic insult. Cancer discovery. 2013;3(11):1254–71.

45. Polkinghorn WR, Parker JS, Lee MX, Kass EM, Spratt DE, Iaquinta PJ, et al. Androgen receptor signaling regulates DNA repair in prostate cancers. Cancer discovery. 2013;3(11):1245–53.

46. O’Neil NJ, Bailey ML, Hieter P. Synthetic lethality and cancer. Nat Rev Genet. 2017;18(10):613–23.

47. Asim M, Tarish F, Zecchini HI, Sanjiv K, Gelali E, Massie CE, et al. Synthetic lethality between androgen receptor signalling and the PARP pathway in prostate cancer. Nat Commun. 2017;8(1):374.

48. Hussain M, Daignault-Newton S, Twardowski PW, Albany C, Stein MN, Kunju LP, et al. Targeting Androgen Receptor and DNA Repair in Metastatic Castration-Resistant Prostate Cancer: Results From NCI 9012. J Clin Oncol. 2018;36(10):991–9.

49. Karanika S, Karantanos T, Li L, Wang J, Park S, Yang G, et al. Targeting DNA Damage Response in Prostate Cancer by Inhibiting Androgen Receptor-CDC6-ATR-Chk1 Signaling. Cell Rep. 2017;18(8):1970–81.

50. Cesare AJ, Hayashi MT, Crabbe L, Karlseder J. The telomere deprotection response is functionally distinct from the genomic DNA damage response. Molecular cell. 2013;51(2):141–55.

